# Genetic basis for antimicrobial resistance in *Escherichia coli* isolated from household water in municipal Ibadan, Nigeria

**DOI:** 10.1101/2024.12.23.630052

**Authors:** Ifeoluwa Akintayo, Jesutofunmi S. Odeyemi, Olumuyiwa S. Alabi, Halimat O. Mohammed, Odion O. Ikhimiukor, Ayorinde O. Afolayan, Nicholas R. Thomson, Iruka N. Okeke

**Author notes:** Address correspondence to Olumuyiwa S. Alabi, and Ifeoluwa Akintayo,. Department of Pharmaceutical Microbiology, Faculty of Pharmacy, University of Ibadan, Ibadan, Oyo State, Nigeria. Institute for Infection Prevention and Control, Medical Center - University of Freiburg, Freiburg, Germany.

## Abstract

*Escherichia coli* serves as an indicator of recent faecal contamination in water, signaling the potential presence of enteric pathogens. The public health impact of *E. coli* in water becomes more significant when strains harbor virulence genes, and may themselves be pathogenic, or antimicrobial resistance genes that can be transferred to pathogens. In this study, we used whole genome sequencing (WGS) to characterize *E. coli* isolated from household water in municipal Ibadan, Nigeria across two seasons. Antimicrobial susceptibility testing was performed on 97 *E. coli* isolates, and their genomes were assembled using SPAdes. Multi-Locus Sequence Types (MLST), virulence genes and plasmid replicons were determined using ABRicate. Antimicrobial resistance genes (ARGs) were detected using AMRFinderplus. Phylogroups and serotypes were determined using ClermonTyper and ECTyper, respectively. A phylogenetic tree was built using RAxML. Of the 97 isolates, 39(40.2%) were multidrug resistant and 13(15.9%) possessed diarrheagenic *E. coli* (DEC) virulence genes. Resistance to individual antibiotics was higher and DEC characteristics more frequent among isolates recovered in the dry season compared to the wet season. Thirty-seven resistance genes belonging to nine antibiotic classes were detected. Majority of the isolates belonged to phylogroup A or B1, 35unique Sequence Types (STs) were detected and there were seven expanded clones of four or more isolates. This study determined that multidrug-resistant *E. coli*, including DEC, were recovered from household water sources in Ibadan. Some isolates were likely derived from point-sources, highlighting the importance of improved water quality management and sanitation in preventing waterborne disease and antimicrobial resistance transmission.

**IMPORTANCE:** Contamination of household drinking water sources by disease-causing microorganisms is a serious public health concern common in African settings. *Escherichia coli*, an indicator of faecal contamination, can also be a reservoir for resistance genes. We have previously reported high frequencies of *E. coli* contamination of household water in municipal Ibadan. In this study we characterized antimicrobial resistance and virulence genes harboured by contaminating isolates. We found potential diarrhoea-causing *E. coli* in water which often carried antimicrobial resistance genes, irrespective of whether or not they were disease causing. Resistance gene carriage was more common among isolates recovered in the dry, as compared to the wet season. This was attributable to resistant lineages of *E. coli* bacteria spreading in the dry season. The work shows the importance of monitoring drinking water in urban African cities like Ibadan and that treating ground water sources may be necessary, particularly in the dry season.

## Introduction

*Escherichia coli* is a well-established microbial indicator of recent faecal contamination of water and is frequently used in water quality assessments globally (1, 2). *E. coli* in water suggests the possibility of the presence of harmful microorganisms that can cause diseases (2, 3). The presence of pathogenic microorganisms in water sources that are designated for domestic purpose in a community is a critical public health situation, particularly for the vulnerable, like children, which requires urgent intervention to disease and outbreaks.

Strains of *E. coli* colonise the gastrointestinal tract (GIT) of humans and animals and most coexist with the host without causing disease unless the immune defense of the host is compromised (4). However, several strains of pathogenic *E. coli* have been reported over the years with the potential to cause mild to severe intestinal or systemic diseases even in immunocompetent hosts (5, 6). These pathogenic *E. coli* strains carry acquired virulence genes (6).

Based on the acquisition of specific virulence factors, host’s tissue tropism and interactions, mechanisms of pathogenesis and clinical syndrome produced, nine different pathotypes of *E. coli* strains that can cause either intestinal or extra-intestinal diseases have been reported (4). These pathotypes are divided into two main groups. Members of the first group are enteropathogenic *E. coli* (EPEC), enterotoxigenic *E. coli* (ETEC), adherent-invasive *E. coli* (AIEC), enterohaemorrhagic *E. coli* (EHEC), enteroaggregative *E. coli* (EAEC), enteroinvasive *E. coli* (EIEC) and diffusely adherent *E. coli* (DAEC) which are termed intestinal pathogenic *E. coli* (InPEC) because of their potential to cause enteric infections, notably diarrhoea. The second group termed extra-intestinal pathogenic *E. coli* (ExPEC), includes uropathogenic *E. coli* (UPEC) and neonatal meningitis *E. coli* (NMEC) (6, 7) which cause urinary tract infections and neonatal meningitis, respectively. ExPEC are also common causes of bacteremia and sepsis and are therefore sometimes referred to as Septicemia associated Pathogenic *E. coli* (SEPEC). Finally, strains carrying very similar virulence gene repertoires to ExPEC may be avian pathogenic *E. coli* (APEC), responsible for severe respiratory infections in poultry. *E. coli* are classified into phylogroups A, B1, B2, C, D, E,F and G as well as clades I to V (8–11). Phylogenetic sub-classifications can be made using seven gene multilocus sequence typing, with the Achtman scheme published by Wirth et al. (12) more commonly employed, cgMLST or whole genome SNP-based approaches.

*E. coli* strains commonly harbor plasmids, transposons, insertion sequences, integron and other mobile genetic elements (MGEs) ( 13–16), which account for their ability to acquire and disseminate virulence and resistance genes within and across clades (17). Indiscriminate use of antibiotics in human and veterinary sectors has selected for antibiotic resistance among commensal and pathogenic enteric bacteria including *E. coli* (18, 19). We have previously shown that *E. coli* is commonly recovered from household water in Ibadan metropolis and isolated 115 strains during a survey of 250 households (20). In this study, we investigated antimicrobial resistance and genotypic characteristics of *Escherichia coli* previously isolated from household water samples in municipal Ibadan, Nigeria and studied their inter-relationships.

## Materials and Methods

### Bacterial isolates

A total of 97 non-duplicated *E. coli* strains out of the 115 previously isolated from contaminated household water samples collected from five municipal local government areas (LGAs) of Ibadan during our earlier wet and dry season surveys (20) were used in this study. *E. coli* ATCC 25922 was used for quality control of biochemical and antimicrobial susceptibility tests.

### Antimicrobial susceptibility testing

Each viable isolate was subjected to antimicrobial susceptibility testing using disc-diffusion method against twelve antibiotics (Oxoid Ltd) belonging to eight classes namely: penicillin (ampicillin - 10µg), tetracycline - 10µg, chloramphenicol - 10µg, trimethoprim - 10µg, trimethoprim-sulphamethoxazole, quinolones (nalidixic acid - 30µg, ciprofloxacin - 5µg), cephalosporins (cefotaxime - 30µg, ceftazidime - 30µg), aminoglycosides (gentamicin - 10µg, Kanamycin - 10µg) and carbapenem (meropenem - 10µg). *E. coli* ATCC25922 was used as control. Each bacterial suspension was prepared by inoculating the bacteria culture from a 24hr Mueller Hinton agar plate into 5mL of sterile normal saline using a sterile inoculating loop and adjusted to 0.5 McFarland standard equivalent. Sterile cotton swabs were used to spread the bacterial suspension over the surface of Mueller Hinton agar (Oxoid Ltd). Standard antibiotic discs were then aseptically placed at equal distance on the inoculated agar plate using an antibiotic disc dispenser (Oxoid). The plates were incubated at 37°C for 24hr before zones of inhibition were measured and recorded. Based on diameters of the zones of inhibition, strains were classified as sensitive (S), intermediate (I) or resistant (R) according to CLSI standard guideline (21) using the WHONET 5.6 software. Isolates that exhibited resistance to three or more classes of antibiotics were classified as multidrug resistant (MDR) strains (22) Multiple antibiotic resistance (MAR) index was calculated by dividing the number of antibiotics to which each of the isolates were resistant to with the total number of the antibiotics tested against each of the isolate (23).

### DNA extraction, library preparation and whole genome sequencing (WGS)

Genomic DNA of the isolates was extracted according to manufacturer’s instructions using Promega extraction kit and quantified using dsDNA Broad Range fluorometric quantification assay, as described in a previous study (24). Libraries were prepared using NEBNext Ultra II FS DNA library kit (New England Biolabs) and Illumina (MiSeq) sequenced.

### Bioinformatic Analyses

*De novo* assembly and speciation were according to the GHRU protocol (available at: https://www.protocols.io/view/ghru-genomic-surveillance-of-antimicrobial-resista-bpn6mmhe). High quality genomes with assemblies <300 contigs, genome sizes ranging from 4.5 −5.5 Mb and an N50 of >25,000 were selected for further analyses (The quality control data can be seen in Supplementary Table S2). The sequence type was determined using Multi-Locus Sequence Typing (MLST) v2.10 (https://github.com/tseemann/mlst), phylogroups determined using ClermonTyper v24.02 (25) and serotypes was done using ECTyper v0.8.19 (26). Genome assemblies were screened for the presence of AMR determinants using AMRFinderplus v.3.10.23 (27). Result from AMRfinderplus was used for the detection of various pathotypes with PathotypeR script (28, 29, 30). Plasmid incompatibility types and virulence genes were determined using the PlasmidFinder (31) and VFDB databases (32) respectively, implemented on ABRicate v.1.0.1 (Available at: https://github.com/tseemann/abricate). The genome assemblies were annotated using Prokka v.1.14.6 (33). The annotated genomes were used as input to characterize the pan-genome using Panaroo v1.2.7 (34). Sequence alignments of the 3092 core genes were concatenated to generate the core genome alignment. Single nucleotide polymorphisms (SNPs) were extracted from the core genome alignment using SNP-sites v.2.5.1 (35). The core SNP alignment was used as input for building a maximum likelihood phylogenetic tree using RAxML v.8.2.12 (36). The phylogenetic trees were visualized and annotated using Figtree v.1.4.4 (Available at: http://tree.bio.ed.ac.uk/software/figtree/) and Interactive Tree of Life (iTOL) v.6.8.1 (37). Genome pairwise SNP data was used to identify genomes clusters characterized by their high sequence similarity in their core genomes, and highlighted clusters.

### Concordance and Statistical Information

Concordance, sensitivity, specificity, true positives, true negatives, false positives and false negatives between phenotypic and genotypic AMR was determined for aminoglycoside, beta-lactam, chloramphenicol, tetracycline, quinolones and trimethropim using R script https://gitlab.com/-/snippets/2050300. Statistical analysis was conducted using R Studio version 2024.04.1+748. Z-test for proportion was conducted to assess statistical difference in the percentage of isolates resistant to each antibiotic between dry and wet seasons. Fisher’s Exact test was used to determine whether (i) the percentage of multidrug-resistant isolates differed significantly across the different water sources and LGAs during dry and wet seasons and (ii) clustering proportion differed significantly between the dry and wet seasons. Chi-squared tests were performed to determine if phylogroups (A and B1) were significantly more likely to be recovered in a particular season, LGAs or water source. We used a *p*-value threshold ≤ 0.05 to determine statistical significance of our test results. To calculate the distance between one water sample point to another defined by latitude and longitude, we used the Haversine formula (38).

## Results

### Phenotypic antimicrobial resistance of *E. coli* from household water

A total of 97 viable *E. coli* isolates (Dry season, n=58; Wet season, n=39) that were previously collected from household water at mapped sites in Ibadan (20) were evaluated. The *E. coli* strains isolated during the dry season exhibited resistance to ampicillin (51.7%, 30/58), nalidixic acid (48.3%, 28/58), tetracycline (46.6%, 27/58), ciprofloxacin (24.1%, 14/58), cefotaxime (19%, 11/58), gentamicin (19%, 11/58), ceftazidime (15.5%, 9/58) and meropenem (5.2%, 3/58). Resistance to antimicrobials was lower in isolates collected during the wet season, except for tetracycline (56.4%, 22/39) (Figure 1). Significant differences in isolate resistance to nalidixic acid (p=0.02), cefotaxime (p=0.01), ciprofloxacin (p=0.01) and ampicillin (p=0.00) were observed between the wet and dry season. Among the 97 *E. coli* recovered, 40.2% exhibited multi-drug resistance (MDR) phenotypes including 28 (48.3%) of the 58 isolated during the dry season and 11(28.2%) of the 39 isolated during the wet season. The seasonal differences in MDR recovery were most marked for well water (Table 1; p=0.02). The median MAR index was 0.15 for well water isolates obtained in the wet season and 0.3 for dry season isolates (Supplementary Table S1). Higher percentages of the MDR isolates were recovered from Ibadan southeast-IBSE (n=9, 60%), and Ibadan southwest-IBSW (n=10, 55.6%), mostly from well water samples (n= 21, 53.8%), during the dry season and lower (0 – 33.3%) during the wet season (Table 1). Fifty-seven (58.8%) of the 97 isolates had MAR index of ≥0.2, comprising of 35(60.3%) and 22 (56.4%), during the dry and wet seasons, respectively (Supplementary Table S1). The MAR index medians were in the range of 0.1-0.2 for isolates from all the LGAs in the wet season. However, in the dry season, they were 0.4 and 0.35 for IBSE and IBSW, respectively, but remained between 0.1-0.2 in the northern IBN, IBNE and IBNW local government areas (Supplementary Table S1).

**FIG. 1:**
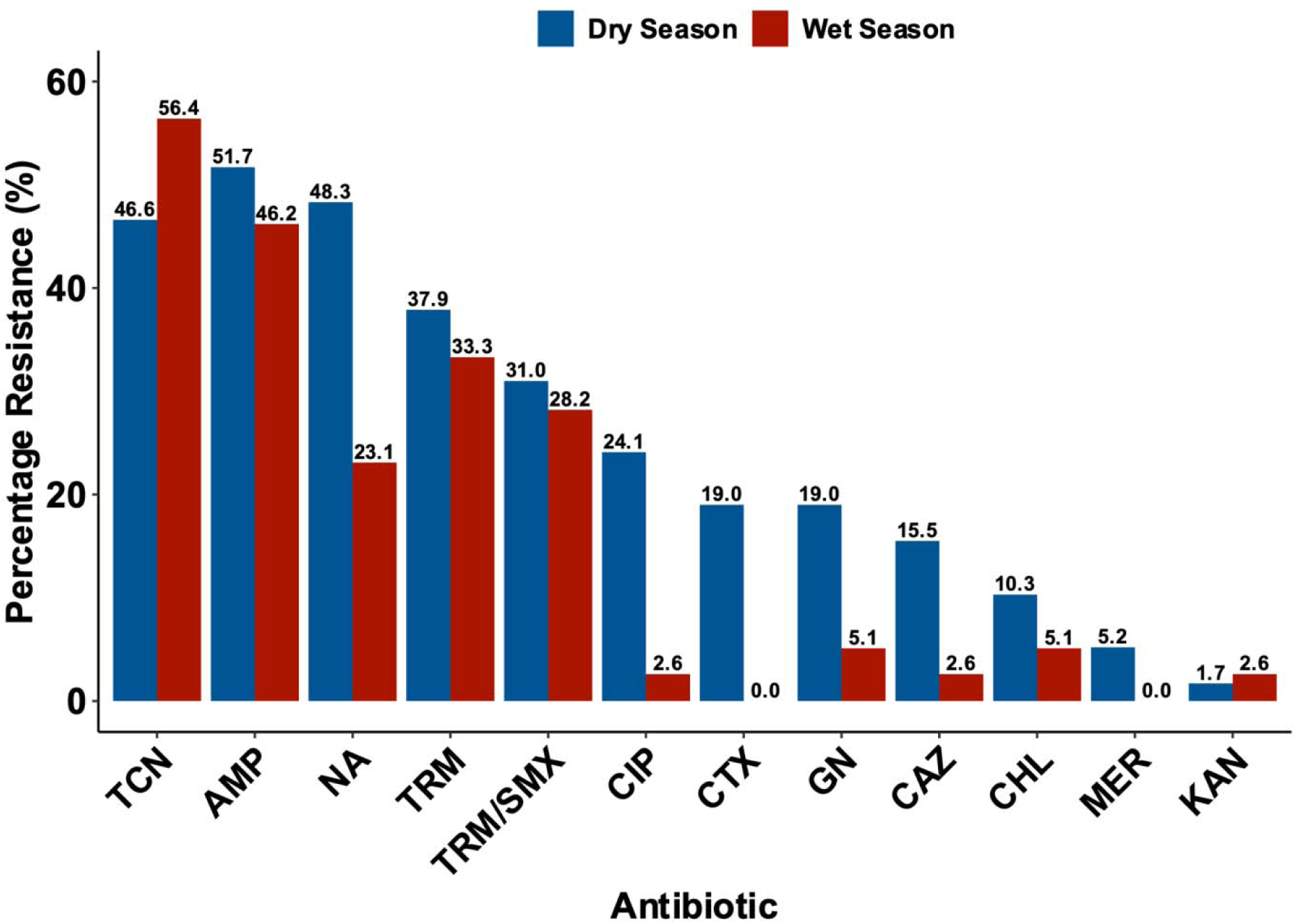
Percentage antibiotic resistance of *E. coli* isolates from water samples during the dry and wet seasons NA – Nalidixic acid, CTX – Cefotaxime, CAZ – Ceftazidime, GN – Gentamicin, MER – Meropenem, CIP – Ciprofloxacin, AMP – Ampicillin, KAN – Kanamycin, TCN – Tetracycline, CHL – Chloramphenicol, TRM – Trimethoprim, TRM/SMX – Trimethoprim/Sulphamethoxazole

**Table 1:** Distribution of the multi-drug resistance phenotype among the *E coli* isolates based on the sources of water, local government areas and seasons.

| Source | MDR (N/%) |  | *P*-value |
| --- | --- | --- | --- |
| Water | Dry | Wet |  |
| Well | 21 (53.8) | 7 (24.1) | 0.02 |
| Borehole | 6 (35.3) | 1 (20) | 1 |
| Stored | 1 (50.0%) | 3 (60.0) | 1 |
| LGA |  |  |  |
| IBNE | 7 (46.7%) | 2 (40.0) | 1 |
| IBNW | 0 (0.0%) | 1 (16.7) | - |
| IBN | 2 (22.2%) | 5 (35.7) | 0.6 |
| IBSE | 9 (60.0%) | 3 (33.3) | 0.4 |
| IBSW | 10 (55.6%) | 0 (0.0) | - |
| Total | 28 (48.3%) | 11 (28.2%) | 0.05 |

### Antimicrobial Resistance Determinants and Plasmid Replicons

A total of 82 (84.5%) of the 97 isolates [Dry season (n = 52) and Wet season (n = 30) isolates] met the quality control threshold for high quality genome and were selected for downstream analysis. The genome sizes ranged from 4.4 Mbp to 5.3 Mbp with percentage G-C of 50.49 – 50.98, and N50 of >53,000. A SNP-based phylogeny of these strains is shown in figure 2. In this study, 37ARGs associated with 16 classes of antibiotics were identified (Fig.3, Supplementary Table S2), including genetic mutations in the resistance determinant regions of core genes (Fig.3, Supplementary Table S2). The most commonly observed were *glpT_E448K*/fosfomycin (n=64, 78%), *tet(A)/*tetracycline (n=39, 46.6%), *sul1*and *sul2*/sulfonamide (n=35, 42.7%), *aph*(*6*)*-Id*/streptomycin (n=34, 41.5%), *bla_TEM-1_*/beta-lactam (n=34, 41.5%), *pmrB_Y358N*/colistin (n=33, 40.2%) and *aph(3”)-Ib*/gentamicin (n=31, 37.8%). Others are *catA1*/chloramphenicol (n=5, 6.1%), *mph(A)*/macrolides (n=5, 6.1%), as well as plasmid-mediated quinolone resistance genes *qnrS1* (n=4, 4.9%), *qnrB19* (n=5, 6.1%), and *qepA4* (n=3, 3.7%) (FIG.3). Resistance based on mutations in the quinolone resistance determinant regions of genes encoding quinolone targets included *gyrA*_S83L (n=26, 31.7%), *gyrA*_D87N (n=13, 15.9%), *parC*_S80I (n=13, 15.9%), *parE*_S458A (n=8, 9.8%), *parC*_S57T (n=3, 3.7%) and *parC*_A56T (n=1, 1.2%).

**FIG.2:**
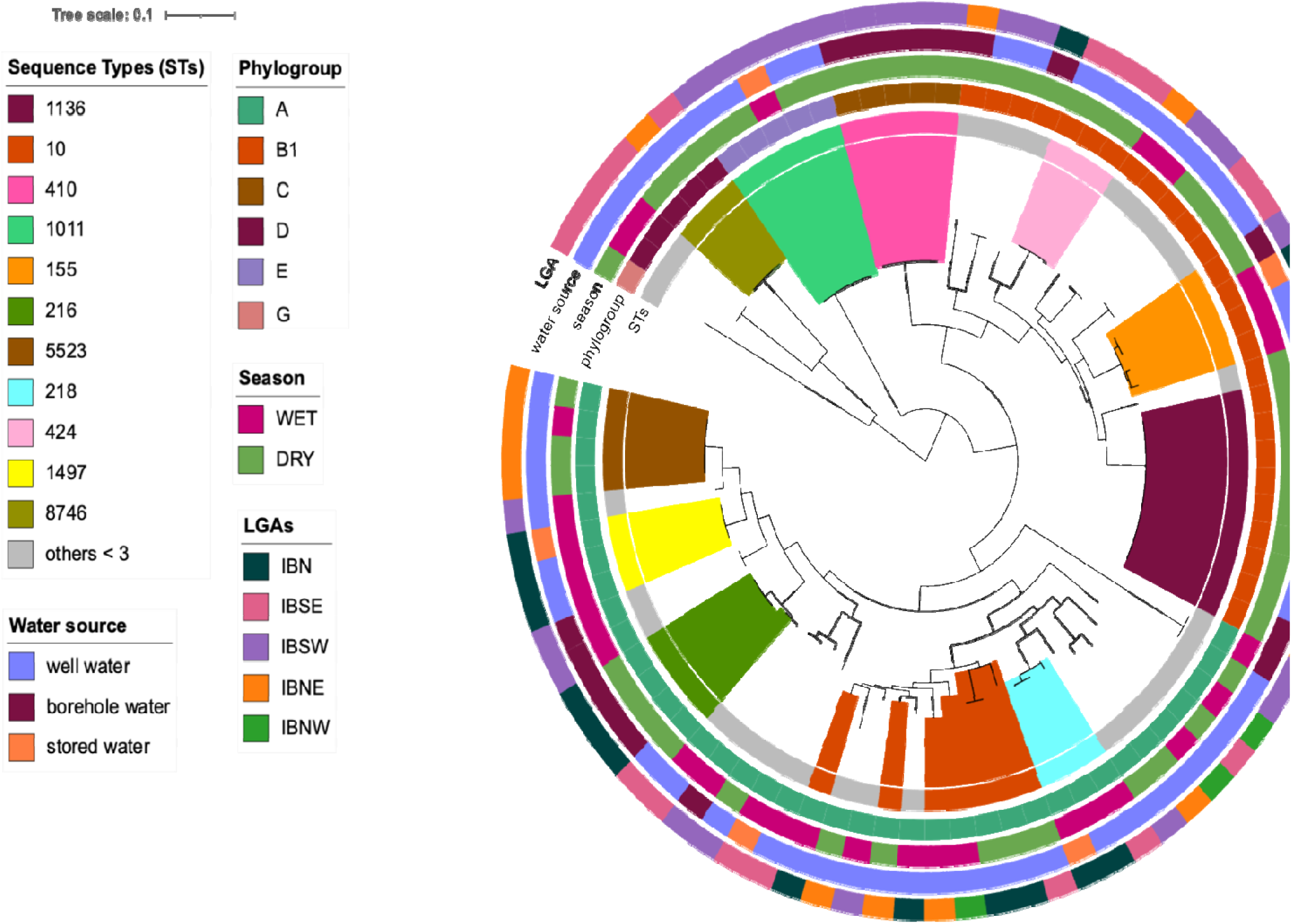
Midpoint rooted phylogenetic tree based on 3,092 core genomes showing the distribution of the sequence types among the water sources and the local government areas during the seasons. (Highlighted are the STs occurring in > 3 isolates. Outer rings showing season, water sources, LGAs and the sequence types. Tree scale represents the number of substitutions per site).

**FIG.3.**
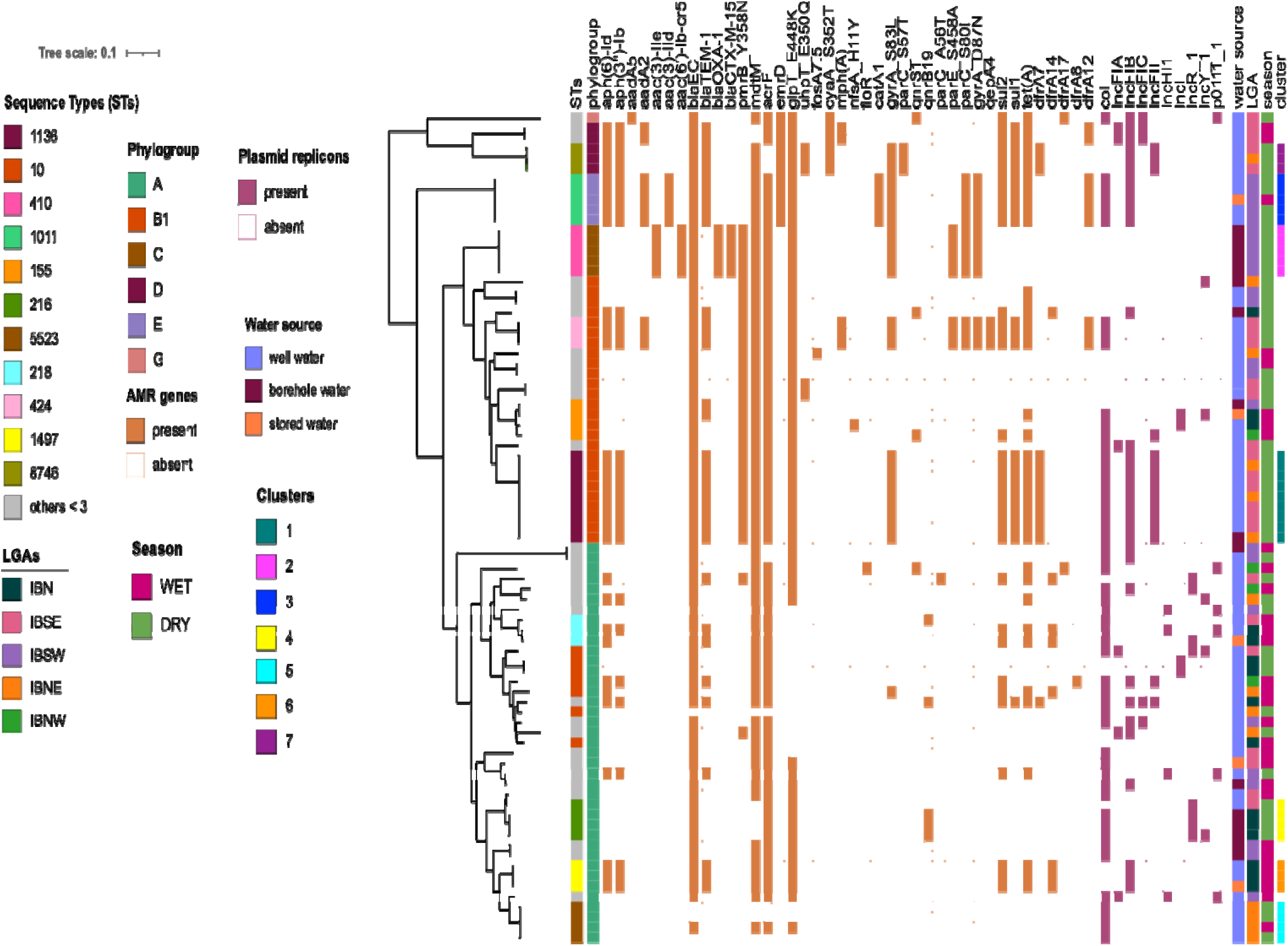
Maximum likelihood phylogenetic tree generated using the single-nucleotide polymorphisms of the core genome of 85 *Escherichia coli* strains characterized by sequence types, clusters, phylogroups, plasmid replicons and the corresponding antimicrobial resistance genes. (Highlighted are the most prevalent ST [occurring in >3 isolates].Tree scale represents the number of substitutions per site).

Forty-one unique plasmid replicons were identified and grouped into *Col* and different *Inc* types (FIG.3). The most prevalent plasmid replicon was *Col*(pHAD28) (n=48, 58.5%), *IncFIB*(AP001918) (n=21, 25.6%), *Col*RNAI (n=14, 17.1%) and *IncFII*(pRSB107) (n=13, 15.9%) (Supplementary Table S2). Other replicons were carried by < 10 representative isolates (Supplementary Table S2). While some of the isolates had multiple replicons (n=53, 64.6%), some others have only one (n=14, 17.1%) or no (n=15, 18.3%) replicon.

A total of 6 classes of antibiotics were tested for concordance between phenotypic resistance and AMR genes. The highest concordance was observed for chloramphenicol (97%) and the lowest was the aminoglycoside (70.6%) (Supplementary Table S3)

### Virulence genes and pathotypes

A total of 204 virulence genes were detected in this study with only 25 virulence genes conserved in all the genomes. These include *acrB, cgsG, csgC,* enterobactin genes *(entA-F and ent S), espl1,* ferrienterobactin precursors and proteins *(fepA-A-D, fepG& fes-1), fur, gndA, ibeB-C, ompA, phoP, resB* and *rpoS*. Membrane transport implicated genes such as those for type VII secretion system, *cfaA,B,C,D,E*(n=>48, >58.5%), type VIII, *csgA,B,CD,E,F,G* (n=77, 93.9%) and siderophore-associated genes, *fepA,B,C,D,E,G* (n=81, 98.8%) were identified. Other virulence genes that were also identified are listed in Supplementary Table S2 and can be explored interactively at: https://microreact.org/project/hcus-ecoli-051224.

Twelves isolates from four water samples in IBNE and IBSE, all recovered during the dry season, were enteroaggregative *Escherichia coli* (EAEC), carrying the *aggR* gene, a plasmid-encoded transcriptional activator, *aap*, dispersin-encoding gene and *aatA-D,* which encode the secretion system for *aap* gene (Supplementary Table S2). EAEC genomes are often enriched for SPATE genes. A total of 10 isolates, including seven EAEC, carried the *sigA* SPATE gene. One water sample from IBNW during the wet season was found to be contaminated with an enterotoxigenic *Escherichia coli* (ETEC) carrying the heat labile enterotoxin *ltcA* gene. However, as many as 47 isolates carried genes encoding colonization factor antigen 1 (*cfa*), including 10 of the EAEC isolates, which are associated with ETEC. No enteropathogenic, enterohemorrhagic or enteroinvasive *E. coli* strains were recovered. A total of 13 (15.9%), 7 (8.5%) and 5 (6.1%) isolates were positive for siderophore genes (*ybt*, *fyuA* and *irp*), pyelonephritis associated pili (*pap*) and alpha haemolysin (*hly*) respectively, marking them as potential extraintestinal pathogens.

### Phylogroups,Sequence Types, and Serotypes of *E. coli* from household water

There are eight principal *E. coli* phylogroups, A, B1, B2, C, D, E, F, and G, all of which have been detected in Ibadan (39). Majority (n=66, 80.5%) of the isolates characterized in this study belonged to phylogroup A(n=40) or B1(n= 26). Strains belonging to *E. coli* phylogroups C (n=5), D (n=5), E (n=5) and G (n=1) were also recovered. While phylogroup B2 strains are common, particularly among pathogens (12), none were detected in this study, although the total number of isolates is small. Phylogroup A isolates were significantly more commonly recovered in the wet season (p=0.001) whilst phylogroup B1 was detected more commonly in the dry season (p=0.04). There were no associations between phylogroups and water sources (well, borehole or stored) or LGAs (p>0.05). Thirty-five different Sequence Types (STs) were identified among the isolates as presented in Supplementary Table S2. The most common STs among the isolates were ST-10 (7), ST-216 and ST-5523 (4) among the phylogroup A strains and ST-1136 (9) and ST-155 (4) among phylogroup B1 strains. ST1136 (n=9, 11%), ST10 (n=7, 8.5%), ST410, ST1011 (n=5, 6.1%) and ST155, ST216 and ST5523 were represented by four genomes (FIG.2). As with their respective phylogroups, some STs were predominantly or only recovered in one season. However, it was observed that isolates belonging to ST10, ST155, ST165, ST1011, ST3696, ST5523, ST12440 and ST12441 were recovered in both seasons (Supplementary Figure 3). Overall, more diversity was seen among isolates recovered in the wet season than in the dry season. A total of 28 (34.1%) STs were comprised of ≤ 3 representative isolates (Supplementary Table S2) these included 18(34.6%, 18/52) of the isolates recovered in the dry season and 20 (66.7%, 20/30) in the wet season.

ST216, ST5523, ST1136 and ST1011/ST410 were the most common in Ibadan north (IBN), Ibadan northeast (IBNE), Ibadan southeast (IBSE) and Ibadan southwest (IBSW) LGAs, with 4, 4, 6 and 5 isolates, respectively (FIG.2). The isolates were distributed into 46 different serotypes with the highest frequency been serotype O59:H19 (n=9), followed by O8:H9 (n=7) and O51:H28 (n=5). All other serotypes were detected at ≤ 4 isolates per serotype (Supplementary Table S2).

### Phylogenetic relatedness and clusters of *E. coli* from household water

The data in figure 4 are juxtaposed against a maximum likelihood phylogeny built from 228,653 SNPs extracted from a 3.38 Mbp sequence alignment of 3,092 shared core genes in the 82 *E. coli* genomes. The data show that the genomes cluster based on STs (FIG.3). Highly-similar clusters were identified based on their pairwise SNP differences, similarity in AMR genes and plasmid replicons. Seven such clusters were identified across LGAs and water sources. The first cluster consisted of nine ST1136 isolates, up to 0 – 6SNPs apart (median 2 SNPs), all belonging to phylogroup B1, which showed identical AMR, plasmid replicons (*Col*(pHAD28), *IncFIB*(AP001918), *IncFII*(pRSB107)) and serotype (O59:H19) with most of them collected from well water (n=8) in two LGAs (IBNE and IBSE). The second cluster includes five ST410 isolates belonging to phylogroup C. All these ST410 isolates were recovered from borehole water in IBSW LGA during dry season with two SNPs differences (FIG.3, Supplementary Table S2). They also show identical AMR profile and serotypes (O8:H9), but no plasmids were detected. The third cluster comprises of five isolates from well water (n=4) (ID5190) recovered during the dry season and stored water (n=1) (ID5120) during the wet season also in LGA (IBSW). These isolates belong to ST1011 and phylogroup E and the locations from which they were recovered are only 2.47 Km apart. They showed genetic similarity differing by only 1 SNP and they also share identical AMR profiles, serotypes (O51:H28) and plasmids (*Col*(pHAD28), *Col*156, *Col*RNAI, *IncFIB*(AP001918)) (FIG3, Supplementary Table S2). The fourth cluster comprises of three isolates from borehole water source recovered during dry season in two water samples from IBN (ID3002 and ID3049). The isolates belong to ST216 and phylogroup A with genetic similarities differing by 3 SNPs. They show identical AMR, serogroup (O86:H9), plasmid replicon (*Inc*R, *Col*(Ye4449) and *Col*(pHAD28)), with an additional plasmid *Inc*Y found in an isolate from sample ID3002. The fifth cluster are four isolates from well water in the same LGA (IBNE) during dry (n=3, ID1073) and wet season (n=1, ID1078). They belong to ST5523 and phylogroup A and the location from which they were recovered are 4.84 Km apart. The isolates are 0-3 SNPs apart, they show similar AMR profile, plasmid replicon (*Col*(pHAD28)) and serogroup (:H21). The sixth cluster comprises of three isolates recovered during the wet season from the same LGA (IBN). Two of the isolates were from well water and one from stored water. They belong to ST1497 and phylogroup A with identical AMR and plasmid replicon (*Inc*FIB(K)). They are 1-4 SNPs apart. The isolates from stored water (ID3056) and well water (ID3078) have identical serogroup (O6:H28). The last cluster, cluster 7, has three isolates recovered during dry season. The isolates were from well water samples from two LGAs, IBNE (n=1) and IBSE (n=2). The isolates showed similarities differing by 0-1 SNP. They belong to ST8746, phylogroup D, serotype O77:H18, with identical AMR and they have the same plasmid replicons (IncFIB(pB171) and IncFII(pRSB107)). All clusters are highlighted in figure 3 and Supplementary Table S2, and can be explored interactively at https://microreact.org/project/hcus-ecoli-clusters-031224. Of the seven clusters, four comprised of isolates recovered only in the dry season, two comprised of isolates recovered in both wet and dry seasons and one cluster comprised of isolates only in wet season. Most clusters contained 3-4 isolates, but one dry season cluster (ST1136) was comprised of nine isolates. Overall, 29 dry season and 4 wet season isolates belonged to one of these clusters (p=0.002, Fisher exact), so that the clusters accounted for lower diversity of isolates in the dry season. And while stored water was an infrequent household water source (n=5) two of the four isolates from clusters in wet season were from stored water. There was at least one cluster in each phylogroup (except G for which there was only one isolate) but five of the seven of the clusters were of MDR strains. As a consequence, 19 cluster isolates were MDR isolates, as compared to 14 of isolates outside a cluster (p=0.01). The median MAR index for isolates inside a cluster was 0.3, while it was 0.2 for those that were not included in a cluster.

## Discussion

We have previously shown that, in the face of failure of the city water system (40, 41), most metropolitan Ibadan residents rely on groundwater for household use (20), a feature that is common in many African settings (42–45). Ground water has been posited as the solution to water access difficulties in both urban and rural Africa (46, 47). However, in areas where sanitation is weak, *E. coli* and other enteric microorganisms can leach into groundwater from animal and human faecal matter (20, 43, 44, 48). Moreover, the warm temperature and high organic matter levels in tropical settings permit enteric organisms to survive and thrive (49, 50). Current attempts at improving Water Sanitation and Hygiene (WASH) in Africa prioritise campaigns against open defecation and establishing standard guidelines by relevant authorities for sinking wells and boreholes within communities. The onus of securing water supplies is left to citizens themselves, who may not take cognizance or be unable to implement key guidelines and thus fall victim to diarrheal diseases and other fecally-transmitted infections (48, 51).

*E. coli* is an important indicator of faecally contaminated water. While the roles of the wild-type *E. coli* strains as important commensals that aid macromolecular digestion and protect the gastrointestinal tract from infections are well known (52), several strains of *E. coli* are pathogenic (53, 54, 55, 56, 57, 58). The role of both non-pathogenic and pathogenic *E. coli* strains as reservoirs for resistance and virulence traits among bacterial species have also been described (3, 23, 59). Therefore *E. coli* can serve as an indicator of the resistance reservoir and provide insight into how resistance genes, and some virulence genes, are spread in communities.

Sammarro et al. (60) recently reported that use of tube wells in Malawi was a risk factor for harboring antimicrobial resistant bacteria. In this study, all the *E. coli* strains isolated from household water at mapped locations across metropolitan Ibadan were antimicrobial susceptibility tested and sequenced. During our isolation, we found and previously reported that 215 of 248 (86.7%) and 146 of 197 (74.1%) household water sources during the dry and wet season respectively, were non-potable across Ibadan metropolis and recovered a total of 115 *E. coli* isolates (20). Ninety-seven of the isolates, studied for this report, exhibited resistance to at least one antibiotic with resistance to tetracycline 49(50.5%), ampicillin 49(50.5%) and nalidixic acid 37 (38.1%) being particularly common. Three of the isolates were resistant to meropenem although no meropenem resistance determinant was found suggesting a possible involvement of unknown resistance determinant to this antibiotic. While *pmrB*Y358N mutation has been reported in both colistin susceptible and resistant *E. coli* with some isolates carrying additional *mcr* gene (61), we cannot conclude about the potential role of this mutation in resistance in our study. Thirty-nine of the isolates (40.2%) were classified as MDR strains. A comparable study in Ghana’s Greater Accra region, which examined a range of water samples, reported that 32% of *E. coli* from well water were MDR (44). Another study in a rural part of Ghana by Kichana et al., (62) reported 48.7% MDR among *E. coli* strains isolated from household drinking water. However, in a previous study by Odonkor and Addo (63), 49.5% MDR strains was reported among *E. coli* isolated from different water sources in Ghana during the dry and rainy seasons. Higher MAR indices in dry season and isolates from IBSE and IBSW could be attributed to several factors such as higher environmental contamination, antibiotic use, hospital practices etc. However, the exact cause remains unclear and requires further investigation.

The number of isolates exhibiting resistance to antibiotics tested in this study was higher among isolates recovered in the dry season. Very few studies examine variations in antimicrobial resistance across season in water for household consumption. However, of those that have multiple studies – including some on groundwater - have reported resistance to be more common among isolates recovered in the dry season, compared to those isolated in wet season (50, 64–66). This is despite the fact that overall contamination by coliforms and *E. coli* is often higher in the rainy season when water tables rise and leaching from proximal unprotected pit latrines or soak-aways may be highest (45, 67–71). While the reason behind the higher prevalence of resistance in dry season is not completely clear, our genomic analysis offers considerable insight. Our data showed that expansion of resistant clones is more frequent in the dry season and in this study, this accounted for the seasonality of resistance, in the largest part. Clusters of identical or near identical isolates were more commonly recovered in the dry season, and more commonly antimicrobial resistant. This is particularly concerning because dry season-amplified clones in this study are remarkably similar to resistant clinical isolates that we have reported previously from Ibadan and elsewhere in South West Nigeria. For example, highly similar multidrug resistant ST410 isolates were recently recovered from patients attending referral hospitals in Ibadan and Osogbo, Nigeria (72) and ST1136 is a little-known EAEC lineage that we recently found to predominate in our setting (39). Thus, groundwater may serve as a reservoir of strains that are virulent as well as resistant (73).

Several virulence genes were identified among the *E. coli* isolates in this study, some of which define pathotypes of virulent *E. coli*. They include: the *aggR* gene, a plasmid-encoded transcriptional activator, *aap* (encodes dispersin which disperses aggregated EAEC) and heat-labile (*lt*) that define ETEC (29) and *aatA-D* genes, which encode the Aap secretion system. Altogether, 12(14.6%, 12/82) isolates were identified as EAEC, all of which were recovered in the dry season. Additionally, colonization factors found in multiple pathotypes, including *csg*A (encodes curli fimbriae for colonisation), *fim* genes (encode type I fimbriae for colonisation, *iucB* (encodes complex siderophore iron receptor), *iutA* (encodes aerobactin synthesis), and *sat* (encodes autotransporter toxin that mediates autophagy) were also detected. The extent to which these and other genes may contribute to survival and transmission in aquatic systems warrants investigation. Recovery of diarrhoeagenic *E. coli* and other pathogens from household water has been reported from other African studies (39, 43, 45, 74) and demonstrates that groundwater may be a vehicle for their spread to susceptible individuals. As many patients with diarrhea do not visit health facilities, and very few health systems have the infrastructure needed to test for diarrhoeagenic *E. coli* in patient stools, the contribution of household water to their transmission is largely unknown.

It is important to note that many of the resistance and virulence genes described among the *E. coli* isolates in this study are typically located on plasmids. Plasmid-borne virulence and resistance genes can be shared among other *E. coli* strains or other bacterial species within the environment. Therefore, the findings in this study have revealed household water sources, particularly well water, as potential reservoirs for multi-drug resistant and pathogenic *E. coli* which thus can pose significant risk to public health.

Eight phylogroups and thousands of STs of *E. coli* have been described. In this study, 35 STs distributed in six phylogroups: A, B1, C, D,E and G, were identified among which STs 1136, 10, 410, 1011, 155, 216 and 5523 were the most commonly encountered. *E. coli* STs 69 and 410 detected in this study have been previously described elsewhere as representing multidrug-resistant extra-intestinal Pathogenic *E. coli* (ExPEC) responsible for invasive urinary tract and blood stream infections (72, 75, 76). The five ST410 in this study were detected in borehole water samples from IBSW and belong to the serotype O8:H9 and phylogroup C. Although no plasmid replicon was identified in the five strains, genes that confer resistance to extended-spectrum beta-lactams (*bla*_OXA-1_, *bla*_CTX-M-15_), aminoglycosides (*aac*(*3*)*-IIe*), quinolones/fluoroquinolones (*aac(6’)-Ib-cr5*, *qepA4*) as well as mutations in the quinolone resistance determinant regions (*gyrA*, *parC* and *parE*) were recorded. Also, the two *E. coli* ST69 detected in this study belong to the serotype O45:H16, phylogroup D and were recovered during the wet season in well water samples from IBSE. These two isolates harbored three plasmid families [IncFIA, IncFIB(AP001918), IncFIC(FII)] and notable resistance genes that mediate resistance against macrolide (*aph*(*6*)*-Id*and*aph(3’’)-Ib*), sulphonamides (*sul1* and*sul2*), trimethoprim (*dfrA12*and *aadA2*), beta-lactams (*bla*_TEM-1_) fosfomycin (*glpT_E448K*), tetracycline (*tet(A)*) and genes coding for multi-drug efflux pumps (*acrF*, *emrD*, *mdtM*) which confer multiple antibiotic resistance phenotype to the strains (77, 78).

More MDR bacteria were recovered from water samples in Ibadan southeast (60%) and southwest (55.6%), mostly from well water samples. It has been reported that water from wells, particularly unprotected wells, can easily be contaminated through human activities and also through close proximity to sewage tanks (79, 80). During the collection of the water samples from which these *E. coli* were obtained, most of the wells were reported to be unprotected (20). Poorly constructed and maintained wells and boreholes can allow contaminant to infiltrate the groundwater and our work identified highly similar isolates recovered from different households showing that, in the absence of effective monitoring and management of water quality, contamination in one area or of one water source can impact the other. Local and national authorities should prioritize raising public awareness on preventive measures by users of non-municipal supplies such as boiling water before use and ensuring human or animal waste are not in close proximity to the water source.

We noted that 5(35.7%) of the 14 strains in dry season clusters carried resistance-conferring SNPs in the QRDRs of quinolone targets and 9(64.3%) of these strains had multiple replicons from broad host range plasmids. Therefore, their amplification and transmission in the dry season has important implications for the epidemiology and ecology of antimicrobial resistance. Rabiu et al., (81) have recently highlighted household water as a useful medium for antimicrobial resistance. Whole genome sequencing is superior for tracking antimicrobial resistance genes and the resistant clones that carry them. However, largely due to reasons of cost, it is rarely employed for this purpose in African settings (82). Our study demonstrates the high level of insight about One Health transmission pathways that can be gained from sequencing isolates from strain sets emerging from carefully designed studies and this is recommended in other instances where the role of specific niches or vehicles in AMR transmission is unclear. Based on the results of this study, we recommend chlorination of household wells, particularly in dry season, to reduce the load or eliminate resistant clones with heightened prevalence in that season. We additionally reiterate our previous finding on non-potability of majority of household water sources in Ibadan and other African settings and the need to improve WASH to avert disease, and as shown in this study, antimicrobial resistance. Moreover, household water and ground waters appear to represent under-appreciated niches for the dissemination of resistant bacteria within and across One Health niches.

One limitation of this study is the change in some household water sampling points during the wet season, many of which were due to consent (20), which prevented us from accessing some previous household water sources directly but instead accessed water point in close proximity. As a result, it was impossible to determine if the same water points that harbored *E. coli* and MDR *E. coli* isolates in the dry season were consistent in the wet season. This also limits our ability to assess whether a particular household water sources is consistently contaminated over both seasons, potentially affecting the interpretation of persistence.

## Conclusion

This study reveals that MDR *E. coli* are commonly found in household water sources in Ibadan. This is particularly true of ground water during the dry season, as well as stored water, which often contain expanded multi-resistant clones. At least some of the isolates are potential pathogens and many have plasmid markers suggesting that at least some of their resistance genes are mobile. The absence of a functional municipal supply and over-reliance on poorly constructed and inadequately maintained wells and boreholes places Ibadan residents at risk of fecally-contaminated water contaminated with *E. coli* that are pathogens and/or could potentially spread resistance to pathogens. Our findings show that safe water and sanitation, while costly to institute needs to be prioritized in our setting. In the interim, raising public awareness on the need to decontaminate household water used for drinking, infant care, and food preparation and access to tools to implement this are important measures that must be taken to mitigate the risks of possible waterborne diseases.

## Acknowledgements

The authors thank field staff recruited for Nigeria’s Severe Typhoid in Africa project’s Health Care Utilization study who collected the water samples from which *E. coli* isolates for this study were obtained. We also thank participants from each household that consented to the study and the staff of the Molecular Laboratory of the Department of Pharmaceutical Microbiology, Faculty of Pharmacy, University of Ibadan (Joy Olorundare, Dr Jeremiah Oloche, Dr David Kwasi) for their technical assistance. We thank Dr Olabisi C Akinlabi for coordinating the quality assurance of the DNA for sequencing and we also thank Rotimi Dada and Erkison E. Odih for helpful discussions.

