## Supplementary figures and images for "Genetic basis for antimicrobial resistance in *Escherichia coli* isolated from household water in municipal Ibadan, Nigeria"

### Supplementary Figure

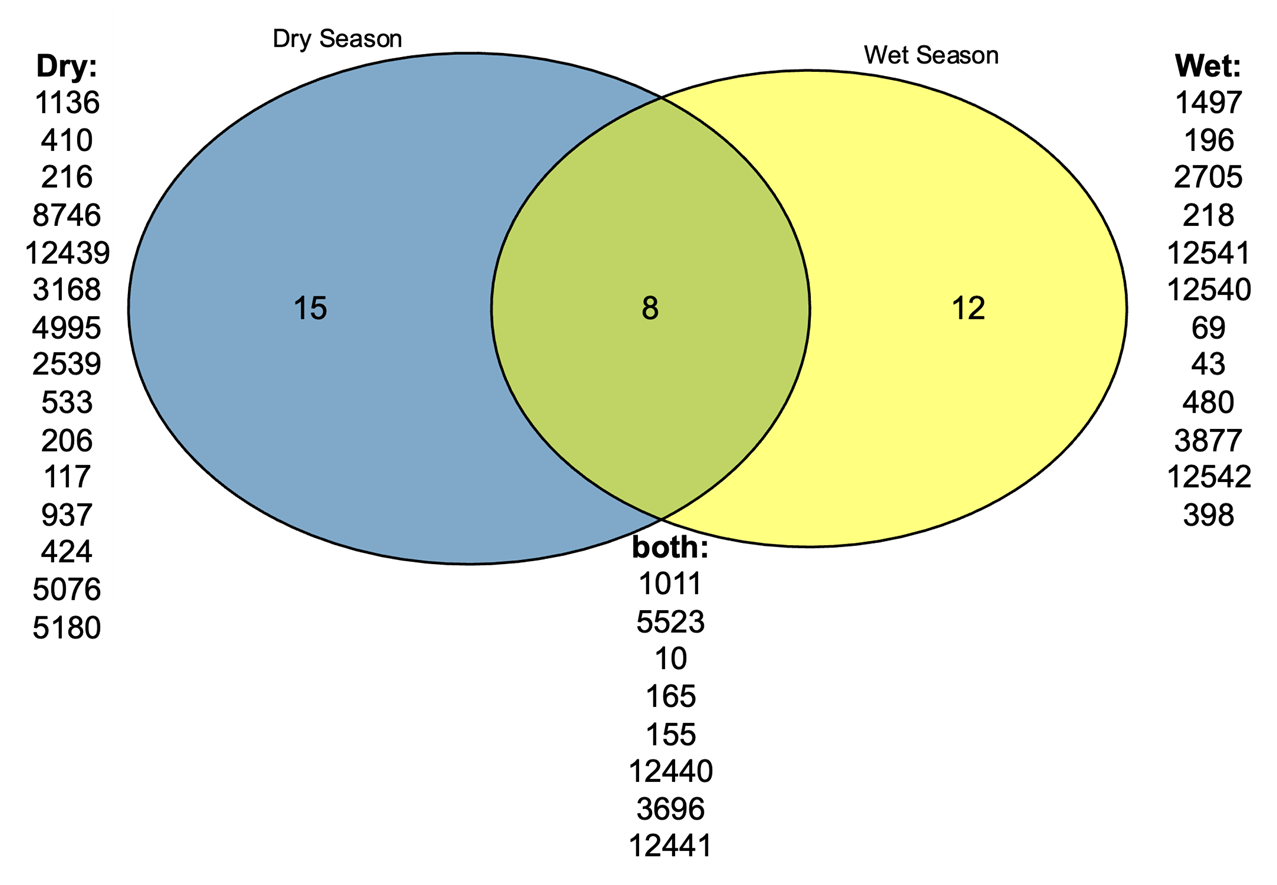

Supplementary Figure 3: Seasonal distribution of the Sequence Types
