## Supplemental Table 1 for "Genetic basis for antimicrobial resistance in *Escherichia coli* isolated from household water in municipal Ibadan, Nigeria"

Supplementary Table S 1: Determination of multiple antibiotic resistance index for isolates from water sources and local government areas

| **Source** | **Dry Season** | | **Wet Season** | |
| --- | --- | --- | --- | --- |
|  | **No of isolate** | **MAR Indices** | **No of isolate** | **MAR indice** |
| Well | 13  4  3  3  6  7  1  4 | 0.0  0.1  0.2  0.3  0.4  0.5  0.6  0.7 | 5  6  3  3  5 | 0.0  0.1  0.2  0.3  0.3 |
| Borehole | 2  4  3  1  2  3 | 0.0  0.1  0.2  0.25  0.4  0.5 | 3  1  1  1  2  1 | 0.0  0.1  0.2  0.3  0.3  0.4 |
| Store tank | 1  1 | 0.1  0.6 | 2  3  3 | 0.1  0.2  0.3 |
| **LGA** |  |  |  |  |
| IBNE | 3  4  1  1  2  2  1  1 | 0.0  0.1  0.2  0.25  0.3  0.4  0.5  0.6 | 2  1  2 | 0.1  0.2  0.3 |
| IBNW | 1 | 0.1 | 1  1  2  2 | 0.0  0.1  0.2  0.3 |
| IBN | 4  3  2 | 0.0  0.2  0.4 | 3  4  2  1  4 | 0.0  0.1  0.2  0.3  0.3 |
| IBSE | 4  2  3  5  1 | 0.0  0.2  0.4  0.5  0.7 | 3  2  1  3 | 0.0  0.1  0.3  0.3 |
| IBSW | 4  4  1  1  4  1  3 | 0.0  0.1  0.3  0.4  0.5  0.6  0.7 | 1  2  1  1 | 0.0  0.2  0.3  0.4 |
